# An in-vivo study of BOLD laminar responses as a function of echo time and static magnetic field strength

**DOI:** 10.1101/2020.07.16.206383

**Authors:** Irati Markuerkiaga, José P. Marques, Lauren J. Bains, David G. Norris

## Abstract

Layer specific functional MRI requires high spatial resolution data. An approach often used for compensating for the poor signal to noise ratio (SNR) associated with small voxel sizes consists of integrating the signal from voxels at a given cortical depth over a patch of cortex. After this integration, physiological noise is expected to be the dominant noise source in the signal. In this context, the sensitivity gain in moving to higher static field strengths is expected to be lower than when thermal noise dominates. In this work, activation profiles in response to the same visual stimulus are compared at 1.5 T, 3 T and 7 T using a multi-echo, gradient echo (GE) FLASH sequence, with a 0.75 mm isotropic voxel size and the cortical integration approach. The results show that after integrating over a patch of cortex between 40 and 100 mm^3^(at 7 T and 1.5 T, respectively), the signal is in the physiological noise dominated regime, and that the obtained activation profiles are similar at the three different field strengths for equivalent echo times. The evolution of the resting-state signal over echo time indicates that BOLD-like noise is the dominant source of physiological noise. Consequently, the functional contrast to noise ratio is not strongly echo-time or field-strength dependent. The results show that compared to 7T, the gold standard, laminar GE-BOLD fMRI at lower field strengths is feasible at the cost of poorer spatial resolution (larger cortical integration extensions) and lower efficiency.

## INTRODUCTION

The human iso-cortex is ~1.5-3 mm thick and is divided into six anatomical layers in most cortical regions. According to the canonical model of neural processing,^1^ layers play a specific role in the processing within a cortical area and in the feed-forward/feedback connections between cortical regions. Therefore, there is a growing interest to acquire high-resolution functional MRI data to measure the layer specific response and understand the interactions between brain regions.

In MRI, high spatial resolution implies a reduced image Signal to Noise Ratio (SNR). Image SNR and temporal SNR (tSNR) increase with the strength of the main magnetic field^2^ Therefore, most layer specific BOLD-fMRI studies in humans in recent years have been performed at 7 T to obtain a higher sensitivity to activation. ^3–13^

Many of these layer specific studies at 7 T further improved the signal to noise ratio (SNR) by integrating the signal over a patch of cortex at a given cortical depth. In an activated region, the larger the number of voxels that are integrated the better the sensitivity to activation will be, provided the noise is not coherent between voxels. That is the case if only thermal noise is present. Once physiological noise sources, *i.e.* those that result in signal modulations unrelated to neuronal changes induced by the functional task, become important, the signal variation will become correlated over voxels and sensitivity will not improve by summation of new voxels. In this physiological noise regime ^2, 14–16^ the sensitivity gain in moving to higher static field strengths will be lower than for the thermal noise regime.

This argumentation would in principle apply to all the forms of functional contrast so far explored for laminar fMRI at 7 T: gradient-echo (GE) BOLD, ^5, 6, 9–12 17–20^ spin-echo based BOLD, ^3, 4, 7, 21^ VASO ^22–24^ and SSFP.^25^ Due to its robustness, high functional sensitivity, and speed of acquisition, GE-BOLD is a widely used approach in layer specific fMRI. An assessment of the potential value of GE-BOLD laminar fMRI at static field strengths below the gold standard of 7 T requires a characterisation of the laminar activation profile as a function of TE, and measurement of the relative sensitivity when data are acquired in the physiological noise regime.

We have previously measured activation profiles and noise characteristics at 7 T. ^8^ Here we repeat the 7 T study and extend this approach to 3 and 1.5 T. In order to perform a comparison based on BOLD signal properties alone, it is important to obtain largely distortion and dropout free images at all field strengths and echo times. This is achieved by measuring with a multi-echo FLASH sequence as the more common EPI image encoding scheme would suffer field-dependent distortion and signal loss. In this manuscript, first the laminar BOLD-dependent signal as a function of TE and relaxation time changes are compared across static magnetic field strengths. These features are independent of the temporal resolution. Subsequently, the noise in the signal, which is a feature that depends on the temporal resolution, is analysed to describe the noise regime for which the functional sensitivity results obtained here apply.

## MATERIALS AND METHODS

Nine healthy volunteers (4 female, 30 *±* 4 years old) were scanned in three separate sessions at 1.5 T, 3 T and 7 T (Avanto, Skyra and Magnetom, respectively, Siemens Healthcare, Erlangen, Germany). The coil configurations used at 1.5 T and 3 T were the vendor-provided 32 Channel head coil for reception and the body coil for transmission. At 7 T, the single channel transmit / 32-channel receive radiofrequency (RF) head coil (Nova Medical Inc., Wilmington, USA) was used. The experimental protocols were approved by the corresponding local ethics committees (“Commissie Mensgebonden Onderzoek regio Arnhem – Nijmegen” for measurements at 1.5T and 3T at the Donders Institute, Nijmegen, the Netherlands. The “Klinisches Ethik-Komitee” of the Universitätsklinikum Essen for measurements at 7T at the Erwin L. Hahn Institute, Essen, Germany). Prior to scanning, all participants gave written informed consent in accordance with the guidelines of the local ethics committees.

### Data acquisition

The session protocol consisted of one high-resolution anatomical scan and one high-resolution functional scan, preceded by a low-resolution functional localizer. The purpose of the latter was to position the thin high-resolution functional slab to cover as much as possible of the primary visual cortex, V1, which would activate in response to the stimulus used.

Whole brain, high-resolution, T1-weighted anatomical scans were acquired using the MPRAGE sequence at 1.5 T and 3 T and the MP2RAGE at 7 T. Functional data for each subject were sampled with the cortical segmentation obtained from the corresponding anatomical scan at 3 T. The relevant parameters of the MPRAGE at 3 T were: TR/TI/TE = 2300/1100/3.15 ms, FA=8°, voxel size=0.8 mm isotropic, GRAPPA factor=2. The anatomical scans at 1.5 T and 7 T were used to calculate the co-registration parameters between the functional scan and the anatomical scan at 3 T. The acquisition parameters for these sequences are given in SI:Table1.

The functional online localizer consisted of acquiring a 2D-EPI sequence while the subject attended to the same stimulus pattern as in the subsequent high-resolution functional scan. Functional localizers were processed online and a map of voxels that responded to the stimulus was obtained. This map was used when positioning the thin slab in the high-resolution functional scan to guarantee that it maximally overlapped with the region responding to the stimulus. The relevant parameters of the functional localizer were the same across field strengths (e.g. 3.0 mm isotropic voxel size, TR=3000 ms, FA=90, 60 measurements), and the TEs were 44 ms, 30 ms, 30 ms at 1.5 T, 3 T and 7 T, respectively. The paradigm used during the functional localizer was 15 seconds of grey screen with a fixation cross, followed by 3 seconds of the same flickering checkerboard as in the functional scan.

The high-resolution functional scan was acquired using a multi-echo 3D-FLASH sequence with parameters (e.g. voxel size and echo spacing/range) matched across fields as far as possible (see Table 1 for sequence parameters). The slab thickness was thinner at 1.5 T and 3 T in order to minimise the acquisition time per volume or volume TR. Compared to 7 T, sequence TRs were longer at the two lower fields. These were limited by the last acquired echo, which was chosen based on literature values of T2* in grey matter (*c.f.* SI table 3a), and lengthened the acquisition. Besides, in-plane acceleration factors achievable with good image quality were lower than at 7 T (R=3 instead of R=4) for the lower fields. ^26, 27^ In consequence, the slab thickness at these fields was reduced in order to reduce the acquisition time per volume and acquire enough volumes for a functional analysis on a reasonable time-scale.

**Table 1:** Relevant acquisition parameters at the three studied fields.

|  | 1.5 T | 3 T | 7 T |
| --- | --- | --- | --- |
| **Functional scan** |  |  |  |
| Sequence | 3D-FLASH | 3D-FLASH | 3D-FLASH |
| Voxel size | 0.75 mm isotropic | 0.75 mm isotropic | 0.75 mm isotropic |
| Slab thickness | 12 mm | 12 mm | 16.5 mm |
| Nr of slices in the slab | 16 | 16 | 22 |
| BW/pixel | 110 Hz | 170Hz | 240Hz |
| TE (10 echoes) | TE1/ΔTE/TE10=<br>7.3/10.6/102.6 ms | TE1/ΔTE/TE10=<br>5.9/8.1/79.0 ms | TE1/ΔTE/TE10=<br>4.8/5.7/56.1 ms |
| Read-out gradient mode | Monopolar | Monopolar | Monopolar |
| TR | 115 ms | 95 ms | 63 ms |
| Volume TR | 160 s | 130 s | 97 s |
| Duration of functional paradigm | 26 min 41 s | 26 min 27 s | 26 min 23 s |
| Number of volumes | 10 | 12 | 18 |
| FA | 25° | 20° | 20° |
| Parallel Imaging (in plane) | 3 (ref lines: 36) | 3 (ref lines: 36) | 4 (ref lines: 24) |
| **Anatomical scan** | MPRAGE with 1 mm isotropic voxel size | MPRAGE with 0.75mm isotropic voxel size | MP2RAGE with 0.75mm isotropic voxel size |

The stimulation paradigm consisted of a circular checkerboard flickering at 7.5 Hz interleaved with a grey screen. Both conditions had a fixation cross in the middle. The duration of the stimulus block was equal to one volume acquisition time at each field (i.e. 160 s, 130 s and 97 s at 1.5 T, 3 T and 7 T respectively). The total acquisition time of the functional protocol was the same across B_0_ (~27 minutes), which allows the comparison of the fMRI performance in the same amount of time (but different numbers of stimulus cycles).

The BOLD response as a function of stimulus duration was studied during piloting to exclude any adaptation effect during the long stimulus ON block in the high-resolution functional acquisition. This was tested at 3 T using one subject and a 2D-EPI sequence, with 3.5 mm isotropic voxel size and TR/TE=2000/30 ms. In this test, the following intra-stimulus patterns were studied: 6 s on – 2 s off, 5 s on-3 s off, 7 s on – 1 s off and 8 s continuously on. Each of the stimulus patterns was repeated 8 times, forming a stimulus ON block with a duration of 1 min 4 s. A stimulus ON block was followed by a grey screen with a fixation cross of 1 min 4 s and the sequence was repeated 3 times. This paradigm was run four times, once per intra-stimulus pattern tested, and the temporal mean and standard deviation of the BOLD response were examined for each of the stimulus-ON patterns. The continuously ON stimulus scored the highest mean BOLD signal with the lowest standard deviation, consequently this was the stimulus mode chosen for the study.

### Data processing

#### Motion correction and co-registration

First, the outer slices at the edges of the 3D functional slab were discarded due to imperfections of the slab profile. The skull was removed using FSL-BET^28^ and all echoes at a given time-point were summed to improve the SNR of the images prior to motion correction. Motion correction was performed in AFNI ^29^ using a linearised weighted least squares algorithm (two passes, zero padded, 7th order polynomial interpolation), realigning all volumes to the first, and high-pass filtered with 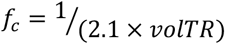. Due to the very thin slab, some of the slices close to the edges suffered from lower or uneven signal intensity in the signal time course due to the interpolation when transforming the volume according to the motion correction parameters. Such slices were removed before further processing. The anatomical volume acquired at 3 T was co-registered to the first functional scan using the boundary based registration method implemented in FreeSurfer^30^.

#### Cortical sampling

The white matter surface was generated in FreeSurfer^10^ and it was expanded by 10% of the cortical depth at each vertex to sample the functional volume (15 sampling points across the cortex: 2 points in white matter, 11 points in the cortex and 2 points in the CSF).^31^ Segmentation performance was visually inspected and corrected in case of major failures. To avoid segmentation inaccuracies less obvious to the naked eye, vertices in V1 areas with an estimated cortical thickness larger than 2.5 mm were discarded from further analysis.

### Feature extraction

Five out of the twenty-seven scans acquired had to be discarded due to within-volume motion artefacts (two at 1.5T and three at 7T), and one scan due to deficient shimming in the occipital lobe. Features were extracted from the remaining twenty-one scans.

#### Signal extraction across the cortex

Depth resolved profiles were obtained by sampling unsmoothed functional volumes using nearest neighbour interpolation with the surface layering obtained in the anatomical scans (see subsection “Cortical sampling” above). All voxels sampled at a given depth were integrated over the corresponding region of interest and the average resting and activation signal over volumes were calculated. The region of interest consisted of:

- only activated voxels for metrics related to activation
- all V1 voxels that fell within the slab for metrics related to noise characteristics

#### Mask of activated voxels

The selection of voxels responding to the stimulus was performed by using FSL-FEAT on the high-resolution data. After summing over all 10 echoes and motion correcting the data, a 3 mm smoothing kernel was applied and voxels that scored z>2.3 were selected for the mask (see Figure 1). Data were smoothed to ensure that the assumptions of random field theory, used to correct for multiple comparisons, were fulfilled, and to improve the sensitivity to activation in acquisitions with a poorer SNR. The surface boundaries obtained in FreeSurfer were overlaid onto the activation maps and vertices falling on activated voxels were identified. This activation map was used to extract the activation related signal for all echo times.

**Figure 1:**
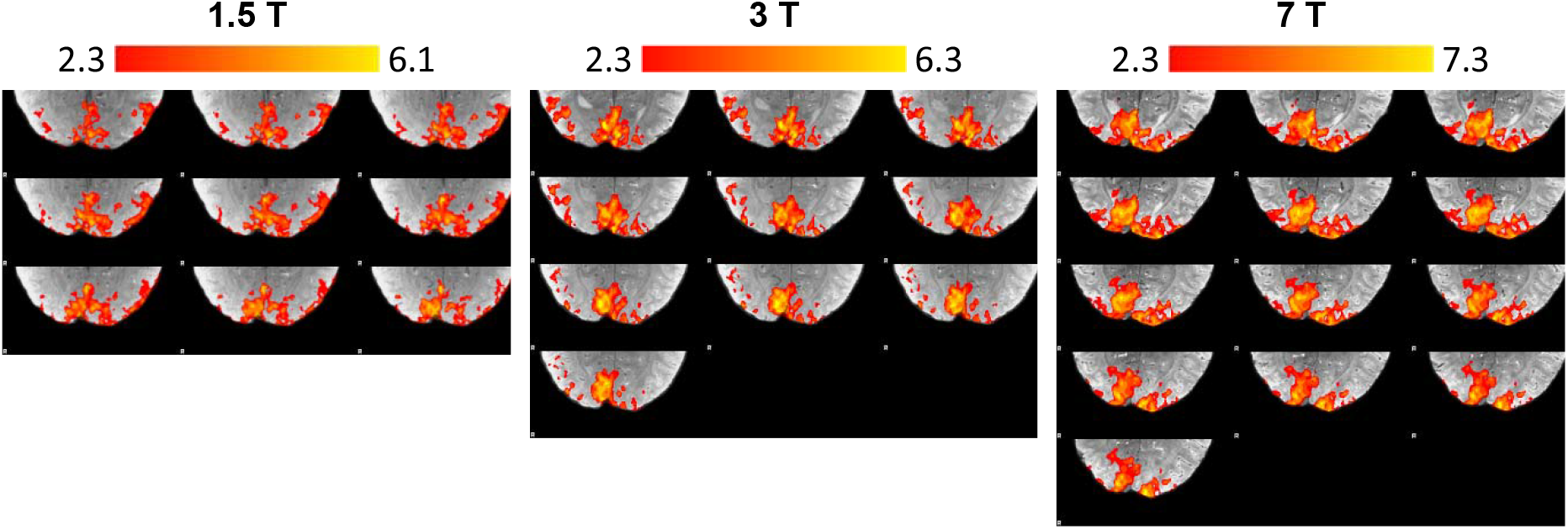
Exemplary activation maps obtained for the same subject at the three different static magnetic field strengths used. Significantly activated voxels are shown in colour. Those in primary visual cortex were used as a mask to extract activation profiles.

#### Feature estimation

R2* was calculated by fitting a mono-exponential function to the signal decay obtained for each cortical depth and volume. The average R2*-values at rest and activation were obtained by averaging over the corresponding state and ΔR2* was obtained as the difference of these two values.

BOLD signal measures were obtained by computing first the average signal in the region of interest per volume and cortical depth and calculating the BOLD signal change and the functional Contrast to Noise Ratio (CNR) per echo time thereafter (a summary of these well-known calculations is given in SI:Table 2).

#### Weisskoff test and physiological noise

The Weisskoff test was described in^32^ as a means of measuring scanner stability. This was performed by measuring, in a phantom, the temporal variance of the mean signal in ROIs of different sizes. The authors in^8^ applied this principle to estimate the minimum extent of the ROI that was needed until physiological noise became the dominant noise source. In order to measure this, an increasing number of voxels from 1 to N, where N is the maximum number of voxels in the area of interest, were averaged, and the temporal standard deviation of the average signal was computed. In the thermal noise dominated regime, this curve is expected to be proportional to 1/√N. If increasing the number of voxels added does not result in a decrease in the standard deviation over time, then the physiological noise dominated regime has been reached.

The Weisskoff-test was carried out by randomly selecting voxels within V1 that were added in each iteration until all voxels belonging to V1 within the slab were part of the ROI. The voxel intensity in an image is in arbitrary units. In order to be able to compare between fields, the standard deviation was normalized using the average signal of all voxels over the complete ROI for TE=T2*_GM_ at each field (Equation 1):

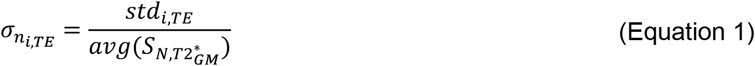

where *i* is the running index that voxels are averaged over before calculating the standard deviation over time, *TE* is the echo time, *S* is the magnitude of the voxel signal and *N* is the number of voxels in the primary visual cortex contained in the slab after motion correction.

## RESULTS

### GE-BOLD signal change across the cortex

ΔS profiles in Figure 2a-c show the characteristic ascending shape of GE-BOLD activation curves across the cortex^8, 10, 24, 33–38^. This ascending pattern is believed to be the result of signal spread through intracortical veins from lower layers^39^ and partial volume effects with pial veins, which will make these profiles steeper close to the cortical surface.

**Figure 2:**
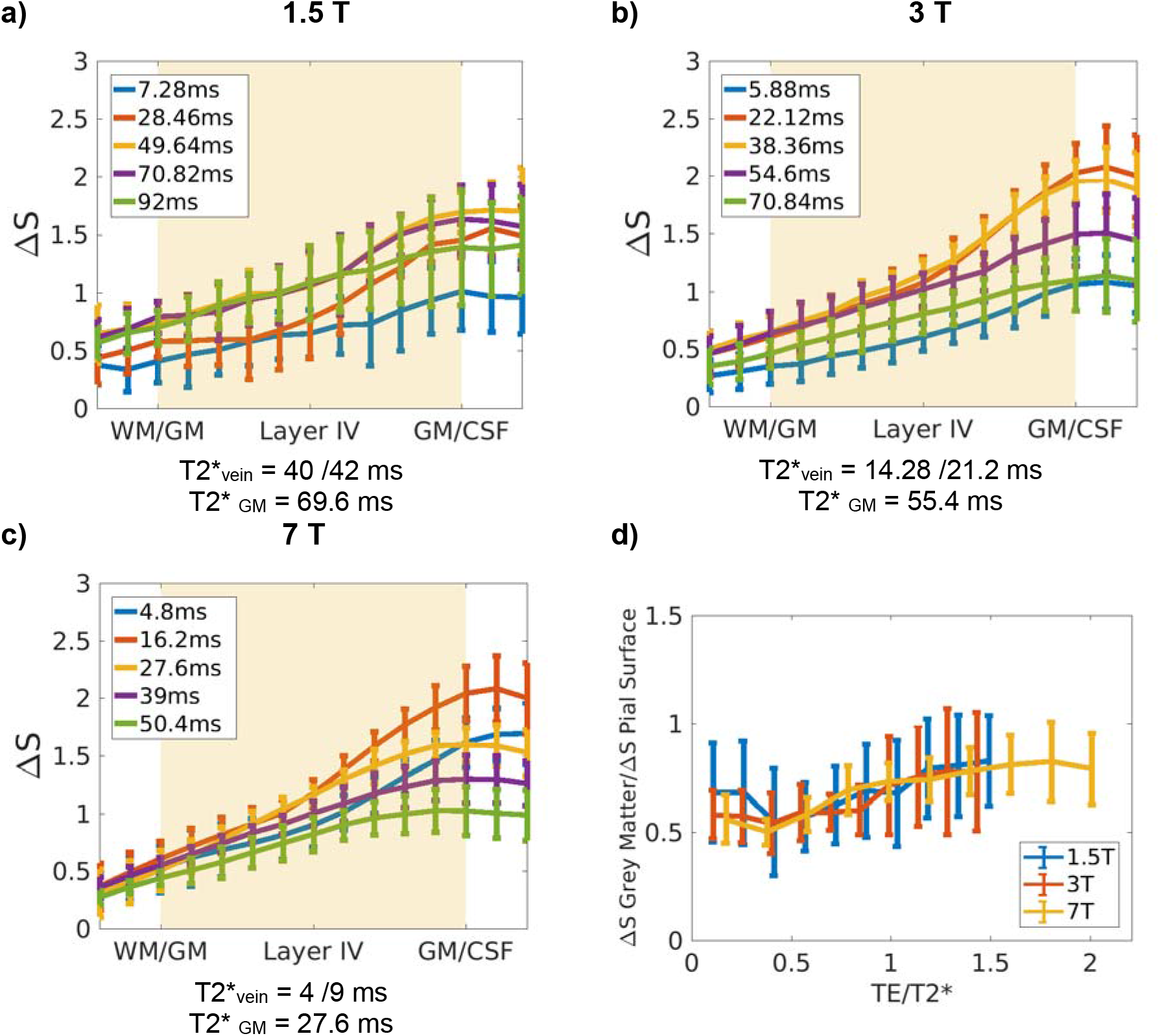
**(a,b,c)** BOLD signal change, ΔS, profiles averaged over subjects for different echo times (5 out of 10 echo times shown for visualization purposes). at 1.5 T, 3 T and 7 T **(a), (b)** and **(c)**, respectively. ΔS is normalised to the mean signal change averaged over depth and TE per subject. The individual profiles have been divided by the mean value over echoes and cortex (hence, one value per profile), so that the average profile is not dominated by differences in the mean response between subjects. **(d)** The ratio between the signal change in the middle of the cortex and on the pial surface across fields. The x-axis shows echo times at each of the field strengths normalized by the corresponding T2* in the middle of the cortex. The 2-way ANOVA significance test shows that only the effect of the echo time was significant (p=0.00). For short echo times (TE1-5), with considerable intravascular contribution, the grey matter to pial ratio is significantly different to that of late echoes (TE8-10), where venous signal has faded away.

The curves in Figure 2a-c have been divided by the mean value of the signal change over echoes and cortex (i.e. one value per profile), so that the average profile is not dominated by differences in the mean response between subjects. This way, it is possible to focus on the differences in the activation profiles across echoes and fields. The profiles obtained at 1.5 T (Figure 2a), which represent to our knowledge the first layer specific functional studies at this field, show qualitatively the same expected behaviour as those obtained at higher fields (Figure 2b and 2c) but with a greater variance. Laminar GE-BOLD profiles at 3 T are less frequent in the literature. Nonetheless the results shown here (Figure 2b) are in line with the ascending profiles for similar stimuli reported previously for this field strength.^38, 40^ The profiles at 7 T (Figure 2c) are similar to those shown in,^8^ only smoother. This is because the processing step used here did not involve manual realignment of the individual profiles to account for inaccurate registration, or segmentation of grey matter boundaries. ^8^

Within a region, the steepness of the profiles depends on the intra- and extravascular signal contribution to the BOLD signal change, which varies with TE and B_0_. For example, at 7 T, the signal on the pial surface is larger for shorter TEs (compare yellow and orange curves). This is because at short echo times, there is still a venous intravascular contribution to the BOLD signal change (a compilation of venous relaxation times at different static field strengths is given in SI:Table 3b). This contribution is greater for large pial vessels on the pial surface than for venous vessels in the grey matter, due to the higher blood volume.

To summarise the data presented in Figure 2a-c, and to allow a simpler comparison between fields, the grey matter to pial ratio of the cortical BOLD signal change was computed (Figure 2d). This was calculated as the ratio between the signal change in the middle of the cortex (which corresponded to the 6^th^ cortical ‘layer’ out of the 11 such layers defined in the cortex by FreeSurfer between white matter and CSF) to the signal change on the pial surface (11^th^ layer in the cortex, which corresponds to the CSF/GM boundary). Note that in the x-axis in this graph TEs -which were different across fields-, have been normalised by the average T2* within the grey matter obtained at that field strength (*c.f.* SI table 3a). For equivalent echo times between fields, the magnitudes of these ratios are similar, around 0.65 for 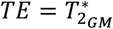. This similarity of the ratios is in line with the simulation results for the ratio of micro- and macrovascular contributions between fields given in.^41^ A 2-way ANOVA test of the pial to grey matter ratio was performed to assess the effects of echo time, field strength, and their interaction. Only the effect of the echo time was significant. For short echo times (TE1-5), with considerable intravascular contribution, the grey matter to pial ratio is significantly different to that of late echoes (TE8-10), where venous signal has faded away.

### Resting R2* and ΔR2* upon activation

Transverse relaxation times at rest and upon activation are shown in Figure 3. The acquisition and analysis approach used in this work (*i.e.* acquiring at submillimetre isotropic resolution and integrating over a given cortical depth) allows for a good separation between cortical signal and signal from the pial surface, as it limits the partial volume effects.

**Figure 3:**
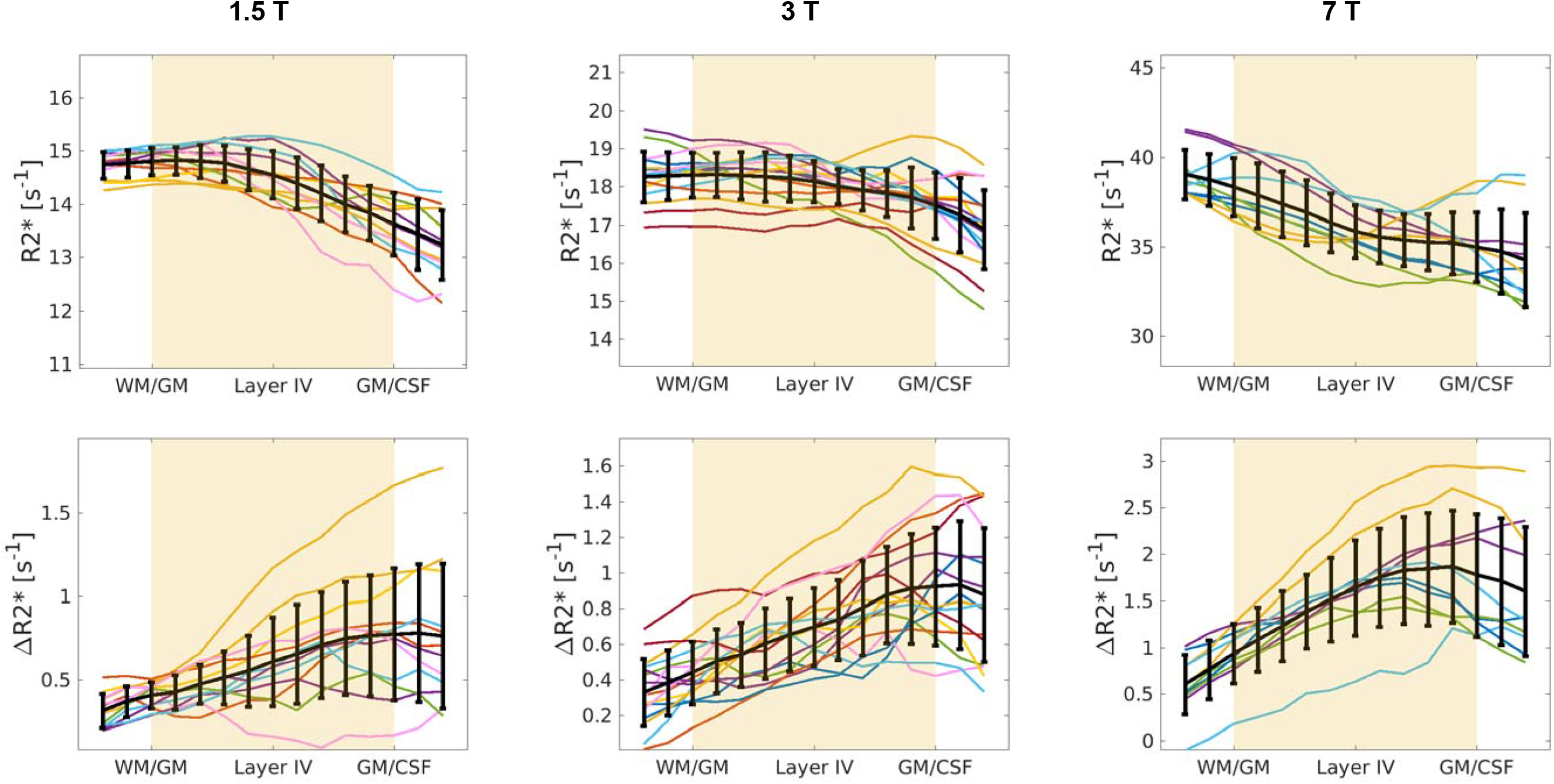
Individual R2* (at rest, upper row) and ΔR2* (during activation, bottom row) profiles for each subject and hemisphere (each colour corresponds to a subject and each line to the profile within a hemisphere). The black line is the mean ± std over subjects, normalized by the average T2* of grey matter at that field.

The mono-exponential signal decay was a good fit for the signal decay within grey matter. The estimated average T2* relaxation times within the cortex (T2*= 70, 55 and 27 ms approximately, see Table 2) are in line with those published previously (as summarised in SI:Table 3a). The average R2* values and profiles obtained at 7 T are similar to those shown in.^42^ The reduction in R2* from the WM to the pial surface (Figure 3a-c) was observed at all field strengths and has been attributed to a combination of reduced myelin and iron.^43^ On the other hand, the variation in transverse relaxation rate upon activation, ΔR2*, showed the opposite behaviour, increasing from the WM to the pial surface, reflecting the increased blood partial volume and oxygenated blood draining towards the cortical surface.

**Table 2:** Comparison of basic activation related results across field strengths.

|  | 1.5 T | 3 T | 7 T |
| --- | --- | --- | --- |
| Number of voxels in the slab after motion correction that fall in the middle of GM | 1199+/-401 | 1140+/-342 | 1514+/-278 |
| % activated voxels in V1 within the slab | 32% | 34% | 41% |
| TE of $\Delta S_{\max}$ on the pial surface | 39.05 ms | 30.24 ms | 10.5 ms |
| TE of $\Delta S_{\max}$ in the middle of the cortex | 60.23 ms | 46.48 ms | 21.9 ms |
| T2* in grey matter at rest | 68.8±2.1 ms | 55.2±1.7 ms | 27.9±1.2 ms |

The voxels that coincide with the pial surface generated in FreeSurfer will in general contain three compartments with very different relaxation times: grey matter, CSF and pial veins. The mono-exponential fit to this signal would over-estimate the change of the relaxation time in the voxel. Unfortunately, it was not possible to fit a multi-compartment relaxation given the noise level in the data, the differences in the relaxation times of the compartments in the voxel, and the range and number of echo times used in this work.

### Temporal fluctuations of the resting signal

Figure 4a-c shows the Weisskoff curves calculated as in Equation 1 are shown in for all subjects and field strengths at TE~T2*_GM_. All Weisskoff curves reach the plateau related to the physiological noise dominated regime after integrating over maximally 250 voxels (about 100 mm^3^), except for one outlier at 1.5 T and 3 T. The differences in the signal to physiological noise ratio between fields (which corresponds to the plateau in the curves) are not statistically significant. A regression analysis between the magnitude of the physiological noise level and the motion parameters from the motion correction algorithm was performed and there was no relationship between the two: scatter plots showed a near random spread of data points (data not shown). Therefore, the variability in the physiological noise level can be related to the variability in the BOLD signal between sessions or/and to other sources of signal fluctuations that cannot be corrected in the post-processing, such as minimal movement during volume acquisition or small differences in the GRAPPA kernel used in each volume. These effects are expected to affect the noise level in the physiological noise dominated regime in a similar way at the three fields. Figure 4d shows the average over subjects of the Weisskoff curves at each field strength, showing how progressive averaging ultimately leads to a plateau in the normalised standard deviation of the resting signal, σ_n_. Visual inspection shows that the point at which the average Weisskoff curve deviates from the noise level in the physiological noise dominated regime (dashed lines), is reached earlier with increasing field, but within 250 voxels at all fields (~100 voxels at 7 T (40 mm^3)^, ~150 voxels at 3 T (60 mm^3^) and ~250 voxels at 1.5 T (100 mm^3^)).

**Figure 4:**
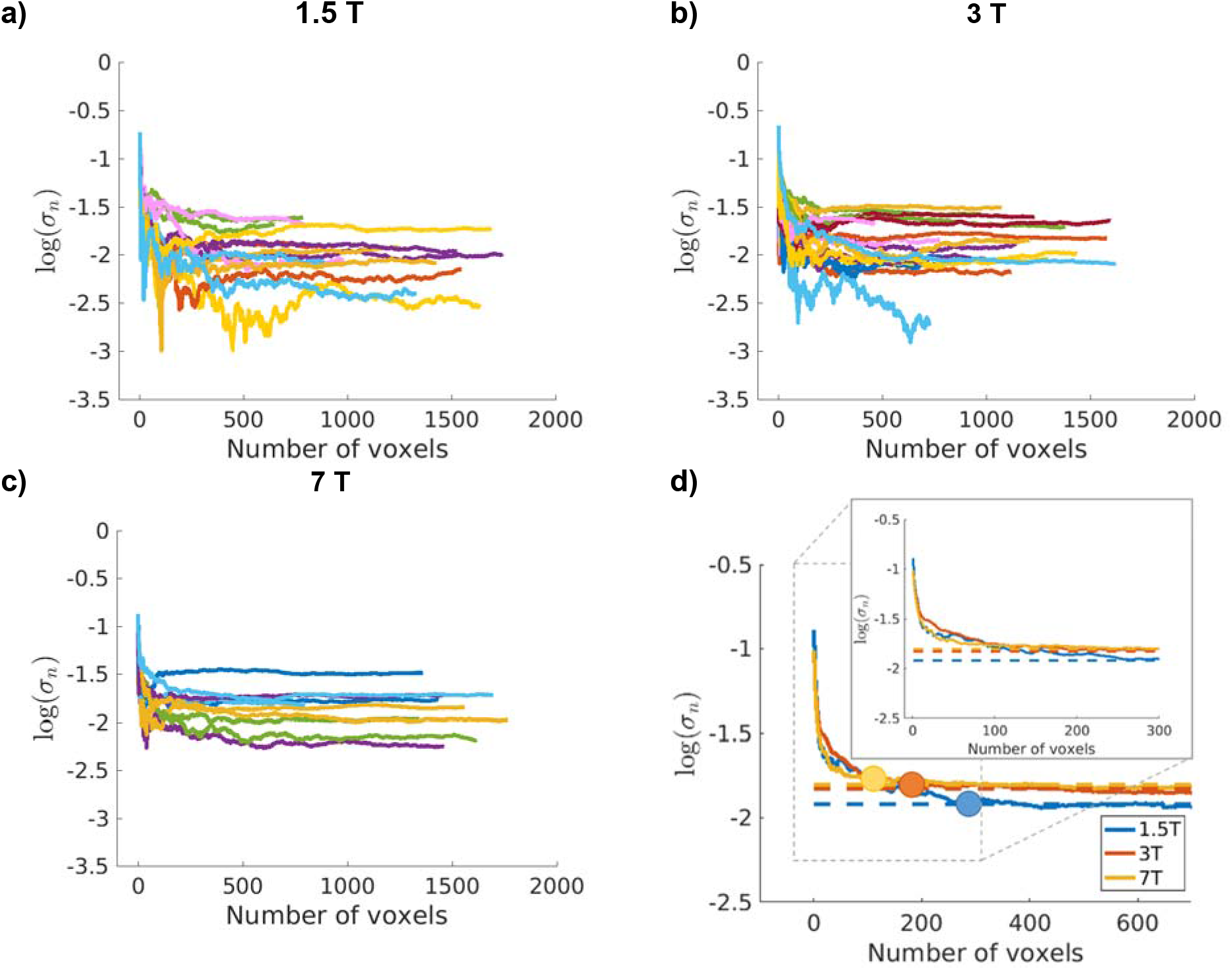
Results of the Weisskoff test: Standard deviation over time of the mean resting signal as a function of the number of voxels averaged (see Equation 1) at 1.5 T, 3 T and 7 T **(a), (b)** and **(c)**, respectively. The voxels added were located in the middle of the cortex and the curves here correspond to TE=T2*_GM_ at each field. Each colour represents a subject, one line per hemisphere. The differences between the physiological noise levels are not significantly different (p=0.49) between field strengths. **(d)** The average of the Weisskoff curves over subjects for the three fields studied. The circles correspond to the point in which the curve deviates from the plateau corresponding to the noise level where the physiological noise dominated regime has been reached (dashed lines with same colour-coding as for static field strength). The inset corresponds to the result for the first 300 voxels (without the circles).

Figure 5 shows how the magnitude of the plateau of the Weisskoff curves in Figure 4 varies as a function of TE. Following the physiological noise model in,^16^ the non-BOLD like physiological noise (e.g. cardiac/respiratory fluctuations) scales with signal strength and will therefore decrease with echo time. The BOLD-like noise (e.g. resting neuronal fluctuations or blood flow changes), on the contrary, will follow the BOLD signal pattern. Figure 5 shows no consistent decrease in noise with TE indicating that BOLD-like physiological noise dominates. Although data have not been corrected for non-BOLD like noise due to the very long volume TRs, the results suggest that BOLD-like noise is the main contributor to physiological noise in this dataset. As a further check, the curves in Figure 5 were compared to those obtained from single voxels, which varied randomly, as expected from a thermal noise dominated distribution.

**Figure 5:**
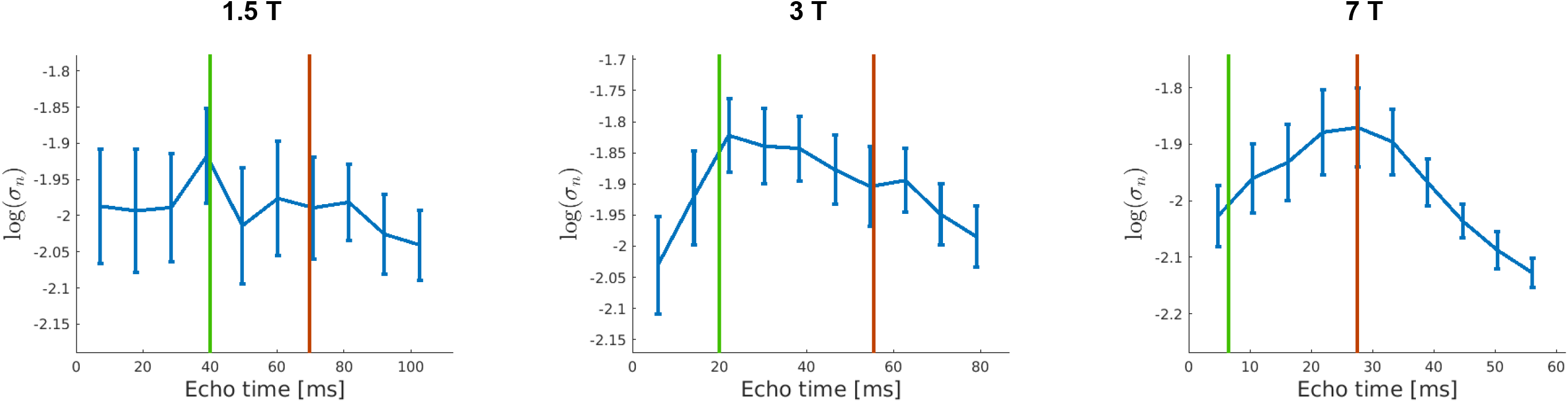
Average physiological noise level over subjects and standard error for the studied echo times in the middle of the cortex at 1.5 T (left), 3 T (middle) and 7 T (right). The green vertical line marks the venous T2* and the red vertical line marks the T2* of tissue at each field. Only the curve at 7T is significantly different from a flat line (p= 0.009).

The 1-way ANOVA test shows that the curves only display a statistically significant dependence of TE at 7 T. However, it is worth mentioning that the profiles for 1.5 T and 3 T show a higher noise level for echo times close to the T2* time of venous blood, marked in green in the figures, than for the echo time corresponding to T2*_GM_, marked in red. This pattern is reversed for the profile at 7 T, which shows a larger magnitude of the fluctuations at around the T2* of tissue. This is in line with the fact that intravascular contribution to the BOLD signal change is larger at 1.5 T and 3 T than at 7 T.^44^

### Functional contrast to noise ratio

Figure 6 shows the functional contrast to noise ratio (CNR) across the cortex obtained after integrating the signal. The CNR is defined as the signal change upon activation (shown in Figures 2a-c) divided by the temporal standard deviation over time at rest. This is considered a measure of the functional sensitivity. As in Figure 2, the profiles are less smooth at 1.5 T, most likely because of poorer co-registration caused by greater difficulties in identifying the GM/WM boundary in the functional scan (refer to Figure 1 for the reduced tissue contrast of the functional scan at 1.5T). The magnitudes of the curves are similar, especially between 3 T and 7 T, as can be expected when physiological noise is the dominant noise source in the signal (see Discussion). The error-bars at 3 T are larger than at 7 T but, if CNR profiles for the first 12 volumes at 7 T are calculated, the size of the error-bars increases (data not shown), indicating that the difference is mainly caused by the higher number of measurements performed per unit time at 7 T. A 2-way ANOVA test with interaction effects shows that within a field and in the middle of the cortex, there is no statistically significant TE-dependence of CNR. The differences in CNR are statistically significant between 1.5 T and 7 T. There is no interaction effect between field strength and echo time. Functional CNR for TE1 to TE3 at 1.5 T are significantly lower than for all TEs at 7 T (p-values = 0.0292 +/− 0.0152).

**Figure 6:**
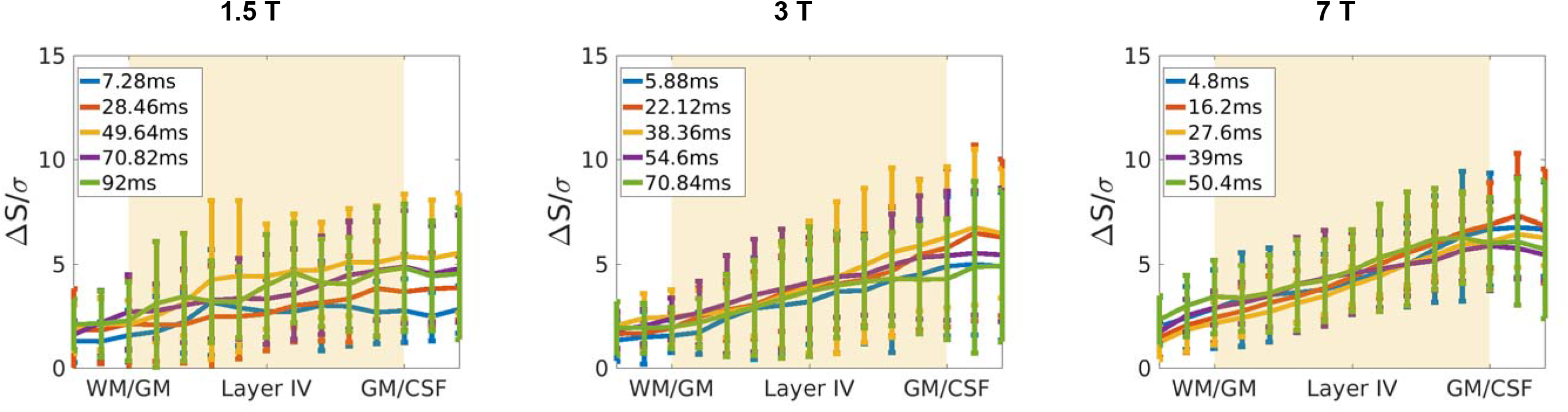
Functional CNR profiles at 1.5 T (left), 3 T (middle) and 7 T (right). The signal change upon activation (ΔS), is divided by the standard deviation over time of the signal in the resting condition, σ. Profiles correspond to average± std over subjects for different echo times (5 out of 10 echo times shown for visualization purposes). The CNR values in the middle of the parenchyma are not significantly different between 3 T and 7 T (p-values ≥ 0.03, threshold corrected for false discovery error rate).

Despite the functional CNR being very similar between 3 T and 7 T, Figure 7a shows that the t-scores obtained at 7 T are higher at all echoes and cortical depths. The temporal resolution is higher at 7 T and more volumes are acquired per unit time, increasing the degrees of freedom of the statistic and therefore the t-score. Within a field strength, the *t*-scores obtained do not vary considerably with TE (Figure 7b). In fact, a 2-way ANOVA test accounting for the interaction effect shows that there is no TE nor interaction effect of TE with field strength. The t-scores obtained at 7 T are significantly different to those obtained at 1.5 T and 3 T.

**Figure 7:**
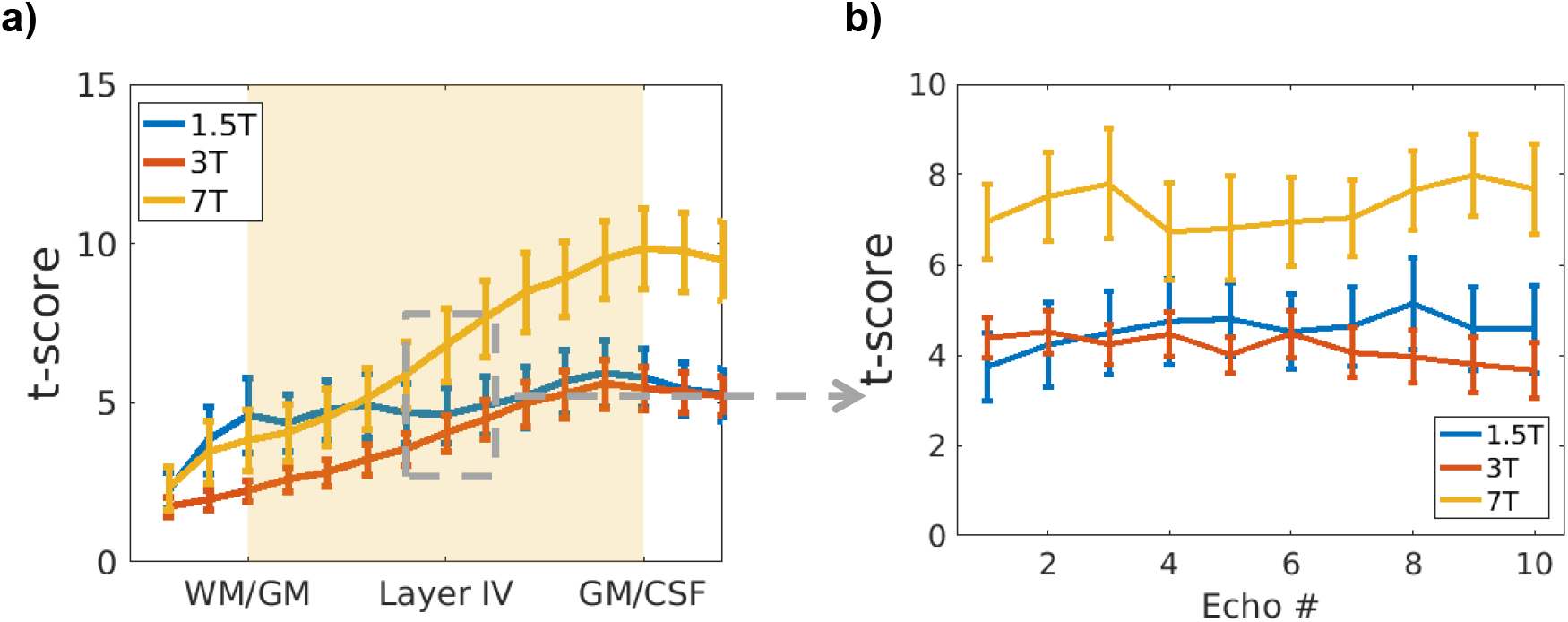
Average over subjects and standard error of **(a)***t*-scores across the cortex for TE=T2*_GM_, and **(b)** t-scores in the middle of the cortex for each of the 10 echoes acquired. Note that the echo times acquired are different for the different static field strengths (see Table 1).

### Mask of activated voxels

Figure 1 shows the activation mask obtained at the three field strengths for a sample subject. The extent of the activation is not identical across field strengths, but they all extend over a representative region of the primary visual cortex. Slight differences in the experimental settings or acquisition (e.g. orientation of the head, placement of the functional slab or visual angle to the screen) are expected to average out after integrating within these masks. A quantitative assessment of the extent of the activation over field strengths and subjects is given in Table 2. The extent of the region responding to the stimulus in the primary visual cortex increases with field strength. As the masks were obtained using the same threshold for the z-score (2.3), the degrees of freedom when performing the statistical inference are higher at 7 T, the z-values will be increased, and more voxels will be above the defined threshold. In addition, the spatial coverage of V1 before and after motion correction was larger at 7 T.

## DISCUSSION

### Scope of this work

This work studies the signal characteristics of layer specific GE-BOLD at 1.5T 3T and 7T, in combination with the commonly used post-processing technique of integrating, or otherwise combining, the depth-dependent signals within a region, to generate a single laminar profile. The GE-BOLD activation pattern depends both on the underlying vasculature, and the neuronal activity. The vascular response can vary over small patches of cortex, among other factors, due to the presence and size of the intracortical veins. As a result, the measured BOLD signal presents some spatial heterogeneity over small patches of cortex, as previously shown.^45^ Our approach, and that of others, has generally been to identify activated regions on the basis of their aggregate response to the task, either by means of an independent activation study, or by smoothing the high resolution data and using this for a non-laminar analysis. Given that we are interested in the characteristics of the ROI as a whole,^5, 18, 46^ an integration approach, or similar, represents the most logical strategy. Small variations caused by vasculature, or indeed gradients in functionality, will then not be accessible, and would require a different approach.

### Activation profiles and relaxation times

GE-BOLD activation profiles are similar for equivalent TEs at 3 T and 7 T. The profiles at 1.5 T show greater variance and a shallower incline going from white matter to CSF for short TEs. The incline is TE-dependent and this fact should be considered, as already mentioned in,^8^ if profiles at different TEs or non-equivalent TEs are compared within or between fields (Figure 2).

The curves in Figure 2d all have a minimum around 0.3-0.4xT2*_GM_ which corresponds roughly to the T2* of venous blood. This shows that the use of shorter echo times increases the pial contribution of the measured signal change. At longer TEs the microvascular contribution to the signal steadily increases for the range of TEs considered in this study.

### Physiological noise dominated regime in the grey matter

The Weisskoff curves in Figure 4 all reach a plateau, the state in which physiological noise is the dominant noise source. The average Weisskoff curves show that this regime is reached after averaging fewer voxels with increasing field strength; around 250, 150 and 100 voxels of 0.75^3^ mm^3^ at 1.5 T, 3 T and 7 T, respectively. An illustration of this for 250 voxels is given in SI Figure 4. Recently, as part of the Human connectome project, researchers were able to parcelate the cortex in 180 region per hemisphere,^47^ resulting in an average surface per parcellated region of 333 mm^2^. Such a region is approximately twice as large as the patch found on this study for 1.5T.

The reduction with increasing field strength in the volume of the cortex that it is required to integrate over to reach the physiological noise dominated regime is of course expected, as the physiological to thermal noise ratio increases with increasing static magnetic field strength.^16^ Most of the improvement in the reduction of the temporal fluctuations is achieved after averaging a small number of voxels: for example, at 1.5 T, around 90% of the total reduction was achieved after averaging over 50 voxels (and sooner for the other two field strengths). These results are in line with the results shown in^14, 15^ in which it is shown that physiological noise dominates even for voxels as small as 3 mm^3^ at 3 T and 7 T when a 32-channel array coil is used.

The integration approach along grey matter used in this work will lead to a higher tSNR as compared to the acquisition of a voxel with an equivalent volume (as long as the voxels have a significant grey matter fraction) as explained in.^48^ An estimation of the increase in tSNR when the integration approach is used, is beyond the scope of this work, as it depends on the spatial extent of the correlation of the physiological noise over the studied region, which is not known.

The physiological noise profiles in Figure 5 show that in this dataset BOLD-like noise is the dominant source of physiological noise. This dominance of BOLD-like noise is probably the result of considering pure grey matter voxels where the BOLD noise should be higher than in white matter or CSF.^49^

### Functional CNR when BOLD-like physiological noise dominates

The time-course CNR determines the functional sensitivity of a sequence and therefore sequence parameters are optimized to maximize this. The optimal TE depends on the dominant source of noise in the signal. If the MR signal in a voxel is accurately characterized by a mono-exponential decay plus some TE-independent noise, i.e., thermal noise, then the highest functional sensitivity or functional CNR will be achieved at TE=T2^(*)^. However, in the presence of physiological noise there will be a noise contribution that is TE dependent.^16^ If the non-BOLD like physiological noise (i.e. mostly the corruption of the signal induced by the heart beat and respiration) dominates, then the maximum of the functional CNR curve will be located at TE > T2 ^(*)^.^50^ If BOLD-like noise (i.e. signal fluctuations related to neural and hemodynamic fluctuations at rest) dominates, then the CNR curve will be flattened and the maximum functional sensitivity will be less TE-dependent.^16, 50^

The CNR profiles across the cortex, see Figure 6, are less steep than the ΔS profiles, especially for shorter TEs. This is because the shape and magnitude of the physiological noise profile as a function of TE follows to some extent that of ΔS, which results in a flattening of the profile and a convergence of the CNR value. In addition, the CNR-values, especially for 3 T and 7 T, are very similar. This implies that when physiological noise dominates, the signal properties of the functional response do not vary strongly with B0.^16, 51^ However, as the temporal resolution in the multi-echo protocols is higher at 7 T, the sequence is more efficient. This results in more degrees of freedom in the statistical analysis, leading to higher t-scores (Figure 7a). As the CNR does not vary considerably between echo times, the t-scores obtained are also very similar between echoes for a given field strength (Figure 7b).

### Laminar fMRI acquisition options when physiological noise dominates

It follows from Figure 7b that, as long as BOLD-like physiological noise dominates, it would be possible to acquire at shorter TEs and improve the temporal resolution of the sequence. Furthermore, note that in the multi-echo acquisition presented in this work, the acquisition at longer TEs increases the TR.

The results in this study also suggest that, if the region of interest spreads over a larger area than the number of voxels required to be in the physiological noise dominated regime at 1.5 T (or 3 T), it should be possible to perform GE-BOLD laminar fMRI studies at field strengths lower than 7 T. Especially at 1.5 T, as the T2* contrast between white matter and grey matter is very poor at this field, it would be advantageous to acquire the functional data with a distortion matched T1-w anatomical scan, as this will considerably ease the segmentation and coregistration of the region^52–54^.

Lastly, for activated regions that are well within the physiological noise regime, it may be possible to increase the spatial resolution, which will potentially allow to differentiate more layers without sacrificing CNR.

### Choice of the acquisition protocol

The primary aim of this study was to characterize BOLD signal profiles across fields and echo times. Therefore, it was important to obtain largely distortion- and dropout-free images to be able to make a comparison based on signal properties alone. Otherwise, the results would have depended on the success of correction-methods in the post-processing. This is why the multi-echo FLASH sequence was used for the functional scan, instead of the EPI approach, more common in functional scans. In addition, the multi-echo implementation in the FLASH sequence allows a more accurate characterisation of the transversal signal decay than would be obtained with multi-echo EPI.

The design of the protocols used was made such that they would be maximally SNR efficient (acquiring during the total TR time), using the same number of echo times, and covering a comparable range of values for TE/T2* of tissue. This was considered the fairest comparison that could be designed across different field strengths.

The Lorentzian line-broadening induced by the relaxation during acquisition is Δf=1/πT2*. In grey matter, this equals 4.6Hz, 5.8Hz and 11.4 Hz, at 1.5 T, 3 T and 7 T, respectively. As the 3D-FLASH sequence was used, the BW per pixels in the PE direction is infinite whereas the BW/pixel-values in the RO direction were 110, 170 and 240Hz/pixel for 1.5 T, 3 T and 7 T, respectively. These values are well above the linewidth, hence they are not expected to have considerable differences in the degree of blurring across fields.

The visual cortex was chosen because it is a well-characterised primary cortical region, in which strong BOLD responses to neuronal activation can be registered. The drawback of studying V1 is that it is rather thin, ~2 mm on average. Therefore, in order to acquire depth resolved signal in V1, the acquisition of submillimetre voxel sizes in the 3 spatial directions was essential (0.75 mm isotropic voxel sizes were used in this study).

All these choices (i.e. distortion free, multi-echo and high-resolution acquisition) provided a high spatial accuracy that came at the cost of temporal resolution (acquisition time per volume over 1 min 30 s at all fields). As the acquisition time per volume varied with field (160 s, 130 s and 97 s at 1.5 T, 3 T and 7 T, respectively), the choice made was to keep the total acquisition time of the functional protocol equal across fields (~27 minutes) and assess the performance in the same amount of time. Because the volume acquisition time at 1.5 T and 3 T was considerably longer, the number of resting volumes at these fields is substantially lower. This results in noisier profiles of the temporal fluctuations of the resting signal, and lower *t-*values for the activation.

### Validity of the results shown here for BOLD fMRI that use a standard temporal resolution

In order to avoid the typical EPI distortion and signal losses that would hinder comparison between fields and echoes, high-resolution functional BOLD data were acquired using the FLASH sequence. As a consequence, the temporal resolution in this work is very poor compared to conventional fMRI studies. It is therefore worth discussing how the results obtained here apply to more common functional GE-BOLD acquisitions based on EPI.

The GE-BOLD profiles as a function of TE and static field strength are independent of the temporal resolution of the sequence. ^55^ reported that for stimuli longer than 20 s the venous compartment dilated, which was not the case for shorter stimuli. The stimulus durations at the different fields used here (i.e. 1 volume TR) are well above that boundary. Therefore, venous contribution across the cortex may be different than those obtained with shorter stimuli. However, this will be the case at the three studied fields (as the stimulus duration is well above 20 s in all cases), so the comparison between fields is still valid.

In terms of the noise behaviour of the resting signal, the results presented here apply to the case in which BOLD-like physiological noise is the dominant source of noise. Depending on the temporal resolution and acquisition scheme (2D/3D) used, the physiological noise level and distribution might be different to that obtained in this work. In addition, the laminar signal obtained in this study is likely to have additional sources of variance, which will appear as “physiological noise”, such as uncorrected small movements within the long volume acquisition times, or the fact that the GRAPPA kernel is calculated for each volume, which might introduce small variations in the time-course.

In order to have an idea of what the physiological noise level is in a more standard fMRI protocol, the Weisskoff test was performed on a functional dataset obtained using a 3D-EPI acquisition at 7 T (other relevant parameters: volume TR ~4 s, 0.9 mm isotropic voxels and TE=22 ms), as shown in SI:Figure 2 in the *Supplementary Information* to this article. The tSNR of the 3D-EPI for the integrated signal in the middle of the cortex was 68, whereas the average at 7 T in the FLASH sequence for the corresponding echo time was 78.9. The profile and physiological noise level obtained for this subject is comparable to those obtained using the 3D-FLASH (none of the datasets have been corrected for physiological noise). As the 3D-EPI was acquired at a single echo time, it is not possible to assess which is the dominant source of physiological noise in this dataset. However, if non-BOLD like physiological noise removal techniques are successfully applied in these datasets, then the dominant noise source would still be BOLD-like physiological noise, and the results regarding functional sensitivity would be unaffected.

Although the behaviour of 3D-EPI with acquisition times above the cardiac and respiratory signal periods might show a similar noise pattern to the data acquired in this study, 2D-EPI acquisitions are likely to show a very different noise pattern when compared to multi-shot acquisitions.^56–58^ Hence, the extent over which it is necessary to integrate might be larger than the one shown in this manuscript.

In summary, this work shows that GE-BOLD signal changes across the cortex following activation are TE-dependent and have comparable activation profiles between fields for equivalent echo times. When BOLD-like physiological noise dominates, which was the case in this study, the Contrast to Noise Ratio (CNR) profiles are largely echo time and field strength independent. This implies that high-resolution laminar fMRI studies can be acquired at TE different to T2*_GM_ and/or at B_0_ < 7 T with comparable CNR. Nonetheless, spatial resolution and the efficiency of the acquisition increases with increasing static magnetic field. Hence, in practice higher field strengths are to be preferred if available.

## ACKNOWLEDGEMENTS

This work was supported by the Initial Training Network, HiMR, funded by the FP7 Marie Curie Actions of the European Commission (FP7-PEOPLE-2012-ITN-316716). The authors would like to thank Paul Gaalman, Lena Schaefer and Stefan Maderwald for technical assistance and training.

Part of this work has been presented in a talk in the 25th ISMRM Annual Meeting in Honolulu.

## Data availability

The data that support the findings of this study are available from the corresponding author upon reasonable request.

## Author information

### Contributions

I.M. contributed to the design of the study, collected, analysed and interpreted the data and wrote the draft manuscript. J.M. contributed to the design of the study and to the interpretation of the data. L.B. contributed to the data collection. DN designed the study, wrote and revised the manuscript, and contributed to the interpretation of the data. All authors read, improved and approved the final manuscript.

