## Supplementary Information for "An in-vivo study of BOLD laminar responses as a function of echo time and static magnetic field strength"

### SI:FIGURES

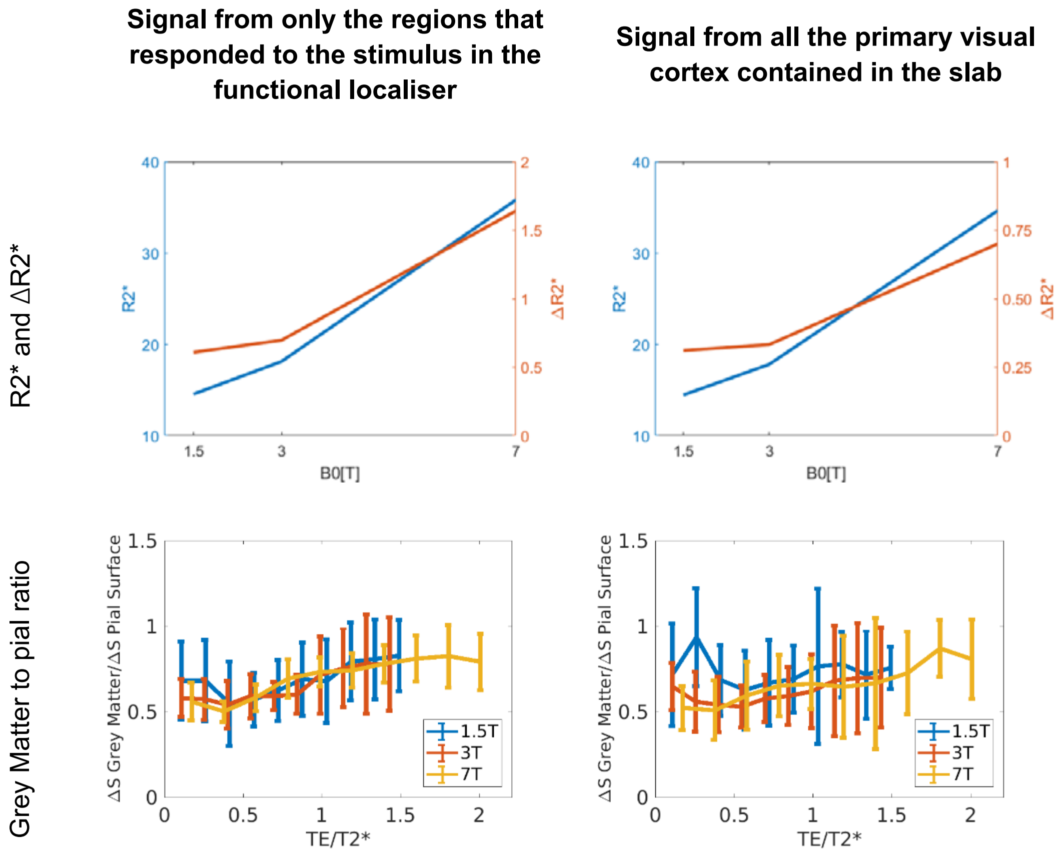


**SI:Figure 1:** Relaxation time values and changes, upper row, and parenchyma to pial ratios, lower row, obtained considering the signal only from the activation mask, left, like in the body of the manuscript and considering signal from all of the primary visual cortex contained in the slab, right.

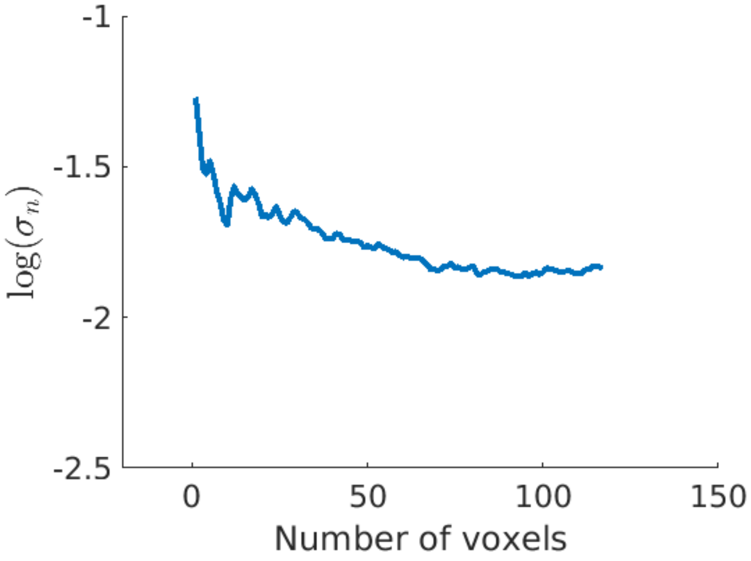


**SI:Figure 2**: Weisskoff test for 3D-EPI with 0.9mm isotropic voxel size and volume TR=3.96 s.

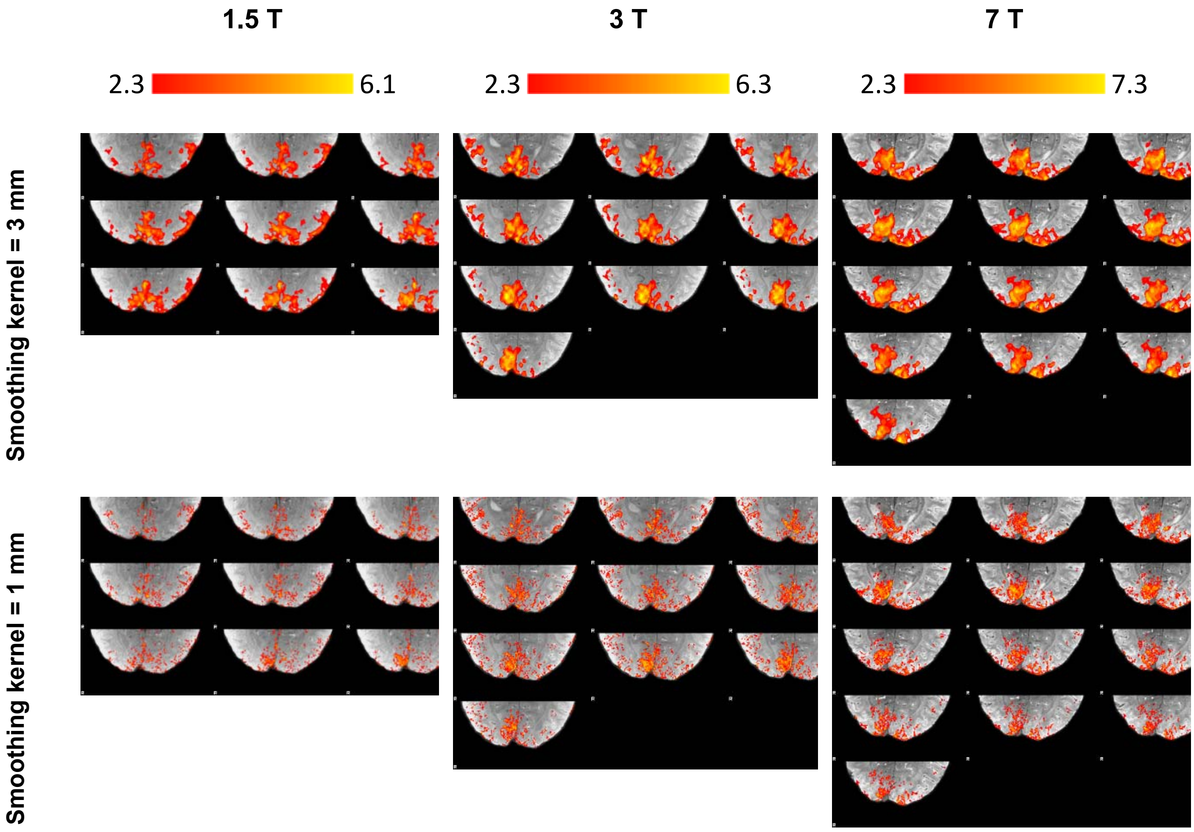


**SI:Figure 3:** A sample of activation maps obtained for the same subject. The upper row shows maps obtained after smoothing with a 3mm Gaussian Kernel, as in the manuscript. The lower row shows high-resolution activation maps, obtained with a 1mm Gaussian Kernel. Data were still smoothed so that the assumptions of random field theory used to correct for multiple comparisons are still valid. The weakly smoothed data show a marginally more restrictive extent of the activation.

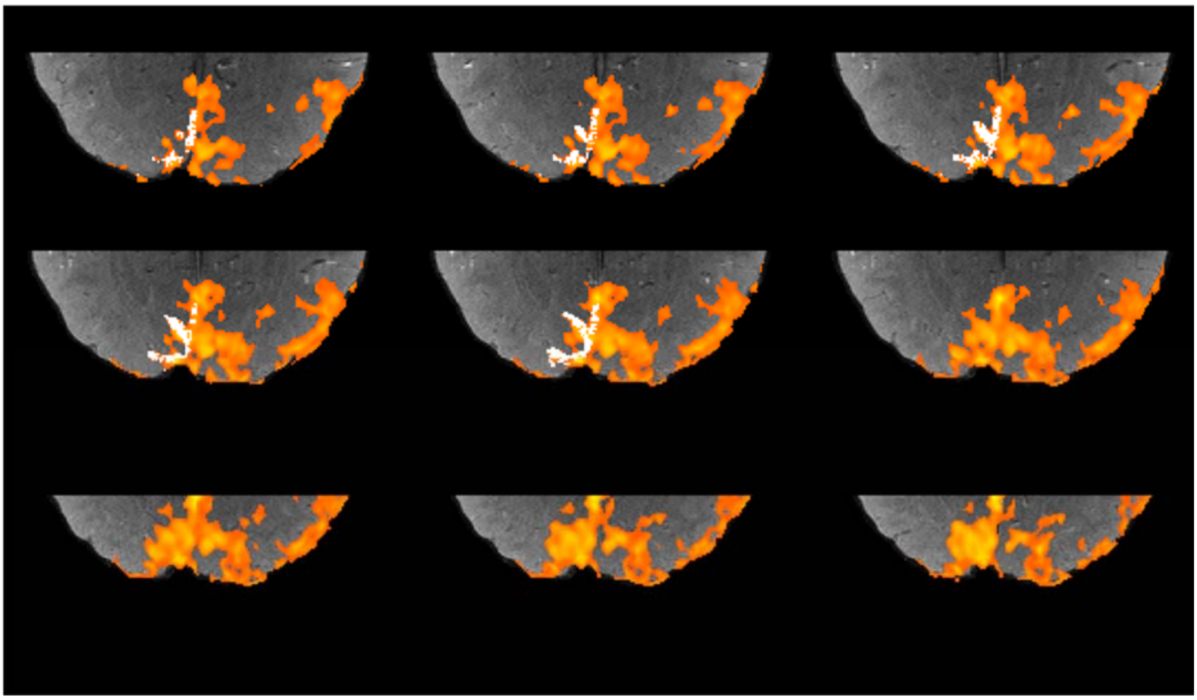

**SI Figure 4:** The area in the occipital region that responded to the stimulus at 1.5T is shown in hot-body scale superimposed on the FLASH image (these are the same data as shown in the left column of Figure 1 in the manuscript). The white overlay shows 250 contiguous voxels chosen arbitrarily from the middle layer of the cortex. For visualisation purposes, also voxels within the grey matter located above and below the mid-cortical voxels are highlighted.

**SI:TABLES**

| | **MPRAGE @ 1.5 T** | **MPRAGE @ 3 T** | **MP2RAGE @ 7 T** |
|---|---|---|---|
| TR | 2300 ms | 2300 ms | 6000 ms |
| TI1/TI2 | 1100/-ms | 1100/- ms | 800 /2700 ms |
| TE | 2.92 ms | 3.15 ms | 3.06 ms |
| FA1/FA2 | 8°/- | 8°/- | 4/5° |
| Voxel size | 1x1x1 $mm^3$ | 0.8x0.8x0.8 $mm^3$ | 0.75x0.75x0.84 $mm^3$ |
| GRAPPA acceleration factor | 3 | 2 | 3 |
| Acquisition time | 4 min 31 s | 6 min 35 s | 9 min 38 s |

**SI:Table 1**: Relevant parameters of the whole brain anatomical scans at each of the field strengths.

| **Name** | **Abbreviation** | **Formula** |
|---|---|---|
| BOLD signal change | ΔS | $\Delta S = \overline{S_{act}} - \overline{S_{rest}}$ |
| Functional contrast to noise ratio | (Functional) CNR | $CNR = (\overline{S_{act}} - \overline{S_{rest}})/std(S_{rest})$ |
| t-score | t | $S = X\beta + \varepsilon$<br>$t = \frac{c'\beta}{\sqrt{Var(\varepsilon)c'(XX')^{-1}c}}$ |

**SI:Table 2**: List of activation related metrics and their functions used in this manuscript.

| Area and paper | Voxel size [$mm^3$] | 1.5 T | 3 T | 4T | 7 T |
|---|---|---|---|---|---|
| **Occipital lobe** [1] | 1x1x3 | 74.9 | 59.7 | | 42.1 |
| **Primary visual cortex**[2] | 1.25x1.25x5 | 69.4 | | 31.7 | |
| **Visual cortex** [3] | 0.75x0.75x5 | | | 41.4+/-5.5 | 25.1+/-3.5 |
| **Average in cortical GM** [4*] | 2.1x2.1x3 | 84.0+/-0.8 | 66.0+/-1.4 | | 33.2+/-1.3 |

**SI:Table 3a:** An overview of $T2^*_{GM}$ values (in ms) reported in the literature.

| Paper | 1.5 T | 3 T | 7 T |
|---|---|---|---|
| **Y=0.6, Hct=0.43**[5] | 40 | 14.28 | 4 |
| **Y = 0.62, Hct=0.44**[6] | | 21.2 | |
| **Y=0.72** [7] | 42+/-2.8 | | |
| **Y = 0.45**[3] | | | 9 |

**SI:Table 3b:** Venous T2* times, in ms) reported in the literature (ex-vivo samples). *Y* refers to the oxygenation of blood and *Hct* to hematocrit.

### *Bibliography (Supplementary Information)*
